# Polycistronic gene expression in the model micro-organism *Ustilago maydis*

**DOI:** 10.1101/2020.05.06.081414

**Authors:** Kira Müntjes, Magnus Philipp, Lisa Hüsemann, Nicole Heucken, Stefanie Weidtkamp-Peters, Kerstin Schipper, Matias D. Zurbriggen, Michael Feldbrügge

**Author notes:** **Correspondence:** Michael Felbrügge.

## Abstract

Eukaryotic microorganisms transcribe monocistronic mRNAs to encode proteins. For synthetic biological approaches like metabolic engineering, precise co-expression of several proteins in space and time is advantageous. A straightforward approach is the application of viral 2A peptides to design synthetic polycistronic mRNAs in eukaryotes. Here, we establish such a system in the well-studied model microorganism Ustilago maydis. Using two fluorescence reporter proteins, we compared the activity of five viral 2A peptides. Their activity was evaluated in vivo using fluorescence microscopy and validated using fluorescence resonance energy transfer (FRET). Activity ranged from 20 to 100% and the best performing 2A peptide was P2A from porcine teschovirus-1. As proof of principle, we followed regulated gene expression efficiently over time and synthesised a tri-cistronic mRNA encoding biosynthetic enzymes to produce mannosylerythritol lipids (MELs). In essence, we evaluated 2A peptides in vivo and demonstrated the applicability of 2A peptide technology for U. maydis in basic and applied science.

## Introduction

In bacteria, gene expression is structured in operons containing polycistronic mRNAs encoding multiple proteins. This has the clear advantage that expression of several proteins can be regulated synchronously using a single promoter and terminator. In eukaryotes, mRNAs are monocistronic and therefore synthesis of each protein can be fine-tuned in space and time. According to the RNA operon model, expression of eukaryotic mRNAs is co-regulated by RNA-binding proteins that determine when and where the corresponding target mRNAs are translated (Keene, 2007). However, for genetic and metabolic engineering it is advantageous to mimic polycistronic mRNAs in eukaryotes for efficient co-regulation of mRNAs in a defined spatio-temporal manner. This circumvents, for example, the multiple uses of identical promoters and terminators, which might reduce overall promoter activity or could interfere with strain generation using homologous recombination (de Felipe et al., 2006;Unkles et al., 2014).

A straightforward approach is the use of viral 2A peptides (de Felipe et al., 2006). These short peptide motifs were first discovered in the foot-and-mouth disease virus (FMDV, F2A peptide) of the *Picornaviridae* virus family (Ryan et al., 1991). Translation of polypeptides containing 2A motifs results in the separation of long viral open reading frames in two units without disassembly of the ribosome (Atkins et al., 2007;Sharma et al., 2012). In a so-called “stop and carry on” mechanism, eukaryotic ribosomes pause at a defined glycine of the characteristic DXEXNPG P motif. The 2A sequence most likely adopts an unfavourable conformation in the exit tunnel, which impairs peptide bond formation between glycine at the P site and the weak nucleophilic imino acid proline at the A site. To overcome ribosomal stalling the translated upstream polypeptide chain with the 2A peptide at its C-terminus is released and translation of the downstream open reading frame carries on using proline as its starting point (Ryan et al., 1991;Atkins et al., 2007}).

This ribosomal mechanism does not function in prokaryotes (Donnelly et al., 1997). However, it is widely distributed in eukaryotes as the activity of 2A peptides has been demonstrated in several organisms ranging from plants and animals to fungi (Halpin et al., 1999;Provost et al., 2007;Kim et al., 2011;Daniels et al., 2014;Unkles et al., 2014;Geier et al., 2015). This allows a broad application of 2A peptides to establish polycistronic gene expression in applied science, for example, in the production of carotenoids in plants (Ha et al., 2010), monoclonal antibodies in animal cell culture (Chng et al., 2015) or natural products in fungi (Ryan et al., 1991;Sharma et al., 2012;Beekwilder et al., 2014;Unkles et al., 2014;Souza-Moreira et al., 2018).

We are studying *Ustilago maydis*, the causative agent of corn smut disease (Kahmann and Kämper, 2004;Brefort et al., 2009). Essential for pathogenicity is a morphological switch from yeast to hyphal growth. The yeast form is non-pathogenic and infected corn has been known as a delicacy in Mexico for centuries, showing *U. maydis* to be a safe for human consumption. This basidiomycete fungus serves as an excellent model system not only for plant pathogenicity, but also for cell and RNA biology (Steinberg and Perez-Martin, 2008;Béthune et al., 2019). During hyphal growth, long-distance transport of mRNAs along microtubules is important. The key factor is the RNA-binding protein Rrm4 that links cargo mRNAs to transport endosomes and orchestrates endosome-coupled translation during transport (König et al., 2009;Baumann et al., 2014;Béthune et al., 2019;Olgeiser et al., 2019). Membrane-coupled translation is a wide spread mechanism and endosome-coupled translation was recently also found in neurons (Cioni et al., 2019;Liao et al., 2019). Besides its role as a model system, *U. maydis* is currently being developed as a production chassis for a wide range of biotechnological relevant compounds. This includes itaconic acid as a chemical platform molecule for biofuels, ustilagic acid and mannosylerythritol lipids (MEL) glycolipids as biosurfactants, and various antibody formats as valuable proteins (Teichmann et al., 2010;Feldbrügge et al., 2013;Sarkari et al., 2014;Terfrüchte et al., 2014;Becker et al., 2019;Stoffels et al., 2020). Furthermore, strains were generated to utilise cellobiose, xylose and polygalacturonic acid as a carbon source in the yeast phase, so that plant cell wall components including pectin can be used as starting point for sustainable production (Geiser et al., 2016;Müller et al., 2018;Stoffels et al., 2020).

*U. maydis* is highly amenable for genetic engineering. Stable strains can be generated by homologous recombination. A comprehensive molecular toolbox, including inducible promoters, fluorescence reporters and epitope tags, is available and plasmid construction is facilitated using Golden Gate, Gibson technology, and Aqua cloning (Gibson et al., 2009;Terfrüchte et al., 2014;Beyer et al., 2015). Here, we add the 2A peptide technology to the growing list of molecular tools in order to further increase the methods spectrum for synthetic biological approaches and biotechnology.

## Results and Discussion

### Establishing a reporter system for screening the activity of 2A peptides

To test the activity of different 2A peptides in *U. maydis,* we designed a bi-cistronic reporter system consisting of the following components (Figure 1A): (i) constitutively active promoter, (ii) upstream ORF encoding a red fluorescent protein, (iii) 2A peptide of interest, (iv) downstream ORF encoding a green fluorescent protein fused to a nuclear localisation signal (NLS), and (v) heterologous transcriptional terminator. Thus, an active 2A peptide would result in increased cytoplasmic red fluorescence while green fluorescence will be located in the nucleus. This enables *in vivo* evaluation of the separation activity (Figure 1A; see below).

**Figure 1.**
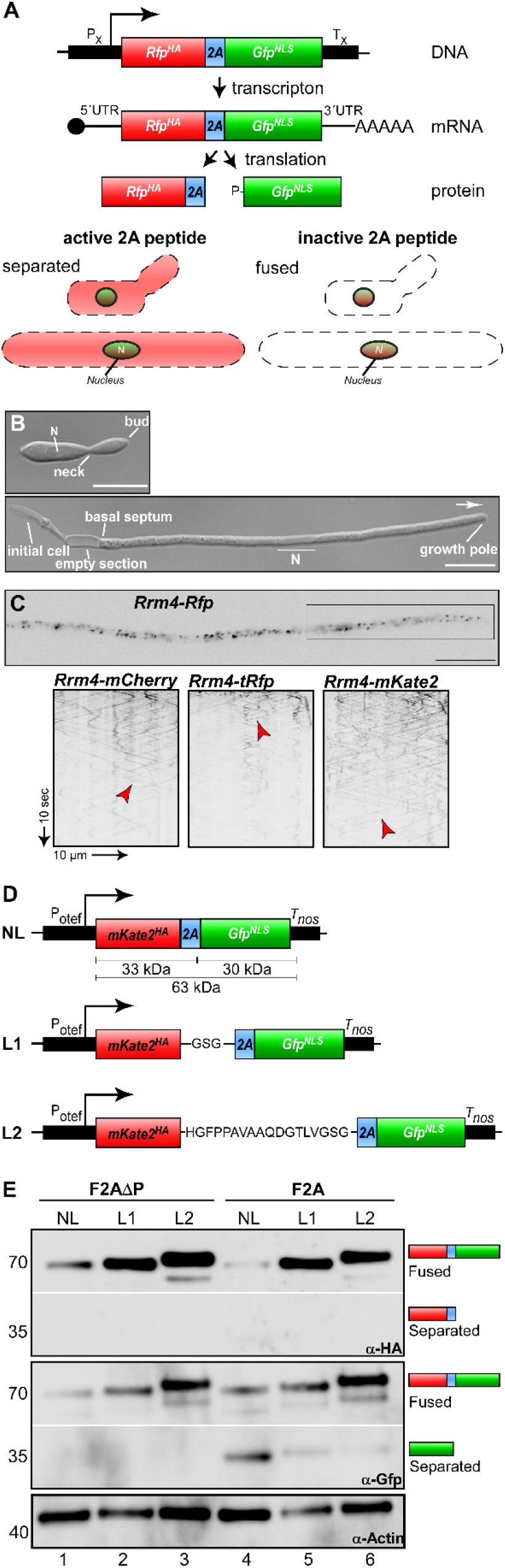
Reporter system for screening the activity of 2A peptides. (**A**) Top: Schematic representation of bi-cistronic reporter: constitutively active promoter (PX), ORF encoding a red fluorescent protein fused to a HA tag (red, RFPHA), 2A peptide of interest (blue) and green fluorescent protein fused to a nuclear localisation sequence (green, GFPNLS), transcriptional terminator (TX). The bi-cistronic mRNA is indicated with 5’ cap structure (filled circle), 5’ and 3’ untranslated region (5’ UTR and 3’ UTR, respectively), and poly(A) tail (AAA). Bottom: Scheme of fluorescent protein localisation within yeast and hyphal cell of U. maydis showing an active 2A peptide (left) or an inactive version (right). (**B**) Growth of laboratory strain AB33 (yeast and hyphae at the top and bottom, respectively). Unipolar growing hypha (6 h.p.i.; growth direction is indicated with an arrow; N, nucleus; scale bar 10 μm). (**C**) Top: Localisation of Rrm4-Rfp within hypha (6 h.p.i.; inverted fluorescence image; scale bar 10 μm). Rectangle indicates region where kymographs were recorded. Bottom: Kymographs of AB33 hyphae (6 h.p.i.) expressing different Rrm4 fusions (arrow length on the left and bottom indicates time and distance). Bidirectional movement is indicated with red arrowheads. (**D**) Schematic representation of linker constructs for 2A peptide analysis. Constitutively active promoter (Potef), mKate2 fused to a HA tag (red), F2A (blue), Gfp fused to NLS (green), transcriptional terminator (Tnos); no linker (NL), GSG linker (L1), 18 aa long linker (L2). (E) Western blot analysis shows ratio of fused to separated proteins (antibodies are given at the bottom, size of marker proteins in kDa at the left).

As a first step we tested different red fluorescent proteins. Currently, the monomeric mCherry protein from *Discosoma* sea anemones is used in *U. maydis* (Baumann et al., 2014). However, the protein exhibits fast photobleaching and its pH stability results in strong fluorescence in vacuoles. This causes difficulties in quantification and localisation of cognate fusion proteins. Therefore, we selected two additional versions, TagRFP and mKate2, both derived from the sea anemone *Entacmaea quadricolor* (Shcherbo et al., 2007;Shcherbo et al., 2009). For evaluation, we generated C-terminal fusions with the RNA-binding protein Rrm4. The correct subcellular localisation of Rrm4 is intensively studied and can easily be scored during hyphal growth because it shuttles on almost all transport endosomes (Figure 1B-C; Baumann et al., 2012;Pohlmann et al., 2015). To generate the fusion proteins, the heterologous ORFs were synthesised according to a context-dependent codon usage that prevents premature poly(A) adenylation of foreign sequences in *U. maydis* (Zarnack et al., 2006;Zhou et al., 2018; http://dicodon-optimization.appspot.com). Corresponding constructs were inserted at the *rrm4* to generate transcriptional fusions loci in the genetic background of AB33 by homologous recombination (see Materials and methods). AB33 is genetically modified to allow an efficient and highly synchronous switch between yeast and hyphal growth by changing the nitrogen source of the medium.

Hyphae expand at the growing tip and insert basal septa resulting in the formation of regularly spaced empty sections (Figure 1B; Brachmann et al., 2001).

All three Rrm4-fusion proteins were fully functional and direct comparison revealed that TagRFP exhibits the highest fluorescence intensity. However, mKate2 is clearly more photostable than the other two fluorescent proteins allowing detailed analyses of subcellular localisation over an extended period of time (Figure 1C). Therefore, we chose mKate2 in our system (Figure 1D-E) and recommend its application in live cell imaging in *U. maydis*.

At the genetic level the respective transcript encoding mKate2^HA^-2A-GFP^NLS^ (eGFP, enhanced version of GFP, Clontech) was expressed under the control of the constitutively active promoter P_otef_ (Figure 1D) and the construct was targeted to the *upp3* locus of the laboratory strain AB33*upp3*Δ by homologous recombination. *upp3* encodes a secreted protease that is dispensable for viability (Sarkari et al., 2014). This targeting strategy was advantageous, since all constructs were positioned at the identical genomic locus and counter-selection (i.e. loss of hygromycin resistance and gain of nourseothricin resistance). This allowed fast pre-screening of homologous recombination events (see Materials and methods). Since it had previously been mentioned that linker sequences influence the expression of open reading frames connected by 2A peptides (Holst et al., 2006;Gao et al., 2012;Souza-Moreira et al., 2018), we tested different linker versions. We compared assemblies without linker (no linker; NL) with those comprising a short GSG linker sequence (L1) and a longer version of 18 amino acids (L2). We started comparing the linker sequences with the canonical F2A sequence from FMDV. As a control we expressed the same sequence without the essential C-terminal proline (F2AΔP; Ryan et al., 1991). As mentioned above all heterologous sequences were designed according to the context-dependent codon usage for *U. maydis* (Zarnack et al., 2006;Zhou et al., 2018; http://dicodon-optimization.appspot.com).

To assess separation efficiency, yeast cells were grown in complete medium and harvested during exponential growth phase. Total protein extracts were tested by Western blot analysis using commercially available α-HA and α-Gfp antibodies (Figure 1E, see Materials and methods). Importantly, comparing constructs without linker to those with linkers L1 or L2, we observed that the absence of a linker resulted in low expression levels (Figure 1E, lane 1 and 4) and the expression level increases with the length of the linker sequence. We observed weak F2A activity for all constructs comprising the F2A peptide (Figure 1E, lane 4-6), but not if the C-terminal proline was deleted (Figure 1E, lane 1-3). Based on these results we chose the linker sequence L2 for further experiments. In essence, we succeeded in setting-up a straightforward reporter system demonstrating that in accordance with earlier reports, proline is essential for activity (Sharma et al., 2012) and linker sequences improve the expression of 2A peptide-containing transcripts (Holst et al., 2006;Gao et al., 2012;Souza-Moreira et al., 2018).

### Screening the Activity of Various 2A Peptides in *U. maydis*

Previously it has been shown that 2A peptides from various viruses exhibited different activities in the ascomycete *Saccharomyces cerevisiae* (Souza-Moreira et al., 2018) as well as in mammalian cells (Kim et al., 2011). In order to identify suitable 2A peptides for *U. maydis*, we selected five versions: four 2A peptides with different separation activities reported from other systems (Table 1; Figure 2A): P2A, Porcine teschovirus-1 (PTV); T2A, *Thosea asigna* virus (TaV); E2A, *Equine rhinitis* A virus (ERAV); F2A, Foot-and-mouth-disease virus (FMDV).

**Figure 2.**
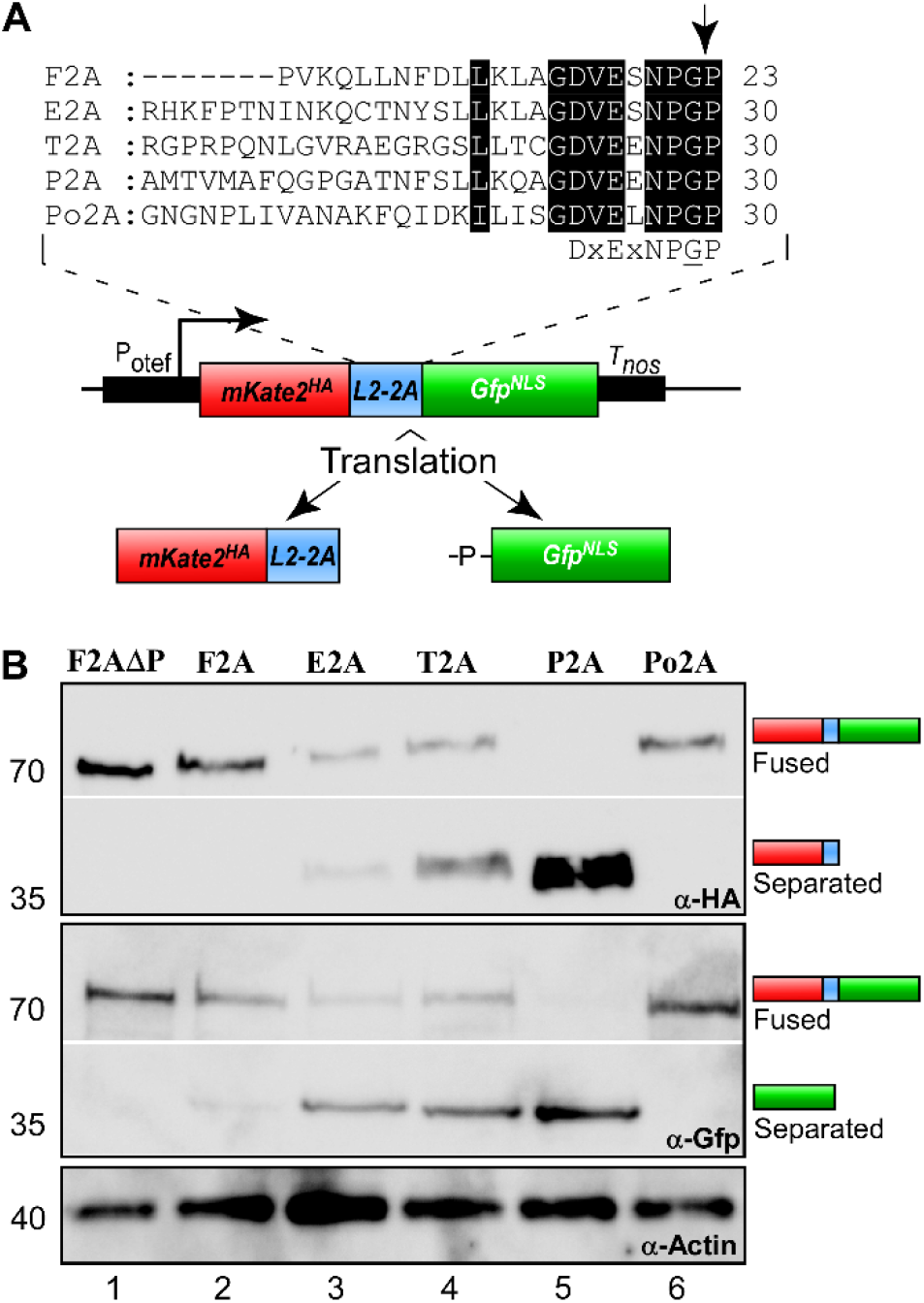
Separation efficiencies of 2A peptides. (**A**) Top: Sequence comparison of 2A peptides (highlighted in black: conserved and important amino acids; arrow indicates point of peptide separation; consensus sequence below). Bottom: Schematic representation of reporter construct and the resulting proteins after translation due to 2A peptide separation: Constitutively active promoter (P_otef_), mKate2 fused to a HA tag (red), L2 linker upstream of the 2A peptide of interest (blue), Gfp fused to NLS (green), transcriptional terminator (T_nos_). (**B**) Western blot analysis showing ratio of fused to separated polypeptides for the different tested 2A peptides (antibodies are given at the bottom, size of marker proteins in kDa at the left).

**Table 1:** Comparison of 2A peptide separation efficiencies in different organisms.

| 2A peptide | Organism | Separation efficiency [%] | Reference |
| --- | --- | --- | --- |
| **F2A** | HEK293T; HT1080; HeLa | ~50 | Kim et al., 2011 |
|  | Zebrafish embryo | ~40 | Kim et al., 2011 |
|  | Mouse liver | ~30 | Kim et al., 2011 |
|  | Silkworm *Bombyx mori* | 91-98 | Wang et al., 2015 |
|  | Drosophila | ~65 | Daniels et al., 2014 |
|  | *Trichoderma reesei* | ~100 | Subramanian et al., 2017 |
|  | *Pichia pastoris* | ~50 | Geier et al., 2015 |
|  | *U. maydis* | 20 | This study |
| **E2A** | HEK293T; HT1080; HeLa | ~60 | Kim et al., 2011 |
|  | Zebrafish embryo | ~60 | Kim et al., 2011 |
|  | Mouse liver | ~50 | Kim et al., 2011 |
|  | Silkworm *Bombyx mori* | 99-100 | Wang et al., 2015 |
|  | Drosophila | ~70 | Daniels et al., 2014 |
|  | *S. cerevisiae* | ~45 | Souza-Moreira et al., 2018 |
|  | *U. maydis* | 60 | This study |
| **T2A** | HEK293T; HT1080; HeLa | ~70 | Kim et al., 2011 |
|  | Zebrafish embryo | ~80 | Kim et al., 2011 |
|  | Mouse liver | ~50 | Kim et al., 2011 |
|  | Silkworm *Bombyx mori* | 90-97 | Wang et al., 2015 |
|  | Drosophila | ~85 | Daniels et al., 2014 |
|  | *S. cerevisiae* | ~55 | Souza-Moreira et al., 2018 |
|  | *U. maydis* | 40 | This study |
| **P2A** | HEK293T; HT1080; HeLa | ~90 | Kim et al., 2011 |
|  | Zebrafish embryo | ~100 | Kim et al., 2011 |
|  | Mouse liver | ~90 | Kim et al., 2011 |
|  | Silkworm *Bombyx mori* | 100 | Wang et al., 2015 |
|  | Drosophila | ~95 | Daniels et al., 2014 |
|  | *S. cerevisiae* | ~85 | Souza-Moreira et al., 2018 |
|  | *Aspergillus nidulans* | ~100 | Unkles et al., 2014 |
|  | *Pichia pastoris* | ~60 | Geier et al., 2015 |
|  | *Aspergillus niger* | ~100 | Schuetze and Meyer, 2017 |
|  | *U. maydis* | 100 | This study |
| **Po2A** | *U. maydis* | 30 | This study |

Fungal organisms shaded in grey

In addition, we chose Po2A from PoRV roavirus C that was not previously studied. Sequences were inserted downstream of the L2 linker (Figure 2A) and respective constructs were again targeted to the *upp3* locus of AB33*upp3*Δ. To assess the different 2A peptide activities, yeast cells were grown in complete medium and total protein extracts analysed by Western blot experiments (see above). The activities were deduced qualitatively from the ratio of fused versus separated forms of mKate2^HA^ and Gfp^NLS^. We observed that F2A and Po2A were hardly active (Figure 2B, lane 2 and 6). There is a clear difference between E2A, T2A and P2A, and the latter showed the highest activity (Figure 2B, lane 3-5).

Next, we studied the separation activity *in vivo*. To this end, we made use of the fact that products localise differently after separation. The separated mKate2^HA^ is expected to localise in the cytoplasm in contrast to GFP^NLS^, which should mainly localise to the nucleus (Figure 1A). A similar set-up with two fluorescence proteins as reporters has successfully been used before to study separation in mammalian cells, silkworm and *S. cerevisiae* (Wang et al., 2015;Liu et al., 2017). In *Aspergillus nidulans*, the activity of 2A peptides was recorded using split Yfp subunits. One half of the fluorescence protein carried an NLS. Hence, expression of the corresponding polycistronic mRNA resulted in nuclear fluorescence (Hoefgen et al., 2018). In our set-up, we fused the NLS to the Gfp reporter (de Felipe and Ryan, 2004;Provost et al., 2007;Kim et al., 2011).

Studying yeast cells revealed that the 2A peptide at the N- and C-terminus did not interfere with the fluorescence of mKate2 or the HA epitope tag. Comparing the 2A peptides we observed that only in the case of P2A we detected strong cytoplasmic red fluorescence, indicating that this 2A peptide seems to exhibit high separation efficiency (Figure 3A). Performing identical analyses in hyphae of *U. maydis* showed the same tendency (Figure 3B, Supplementary Figure S1). Hence, with our set-up we could easily test different stages of the fungal life cycle verifying that there is no developmental regulation of 2A peptide activity. In both cases P2A was the most promising candidate.

**Figure 3.**
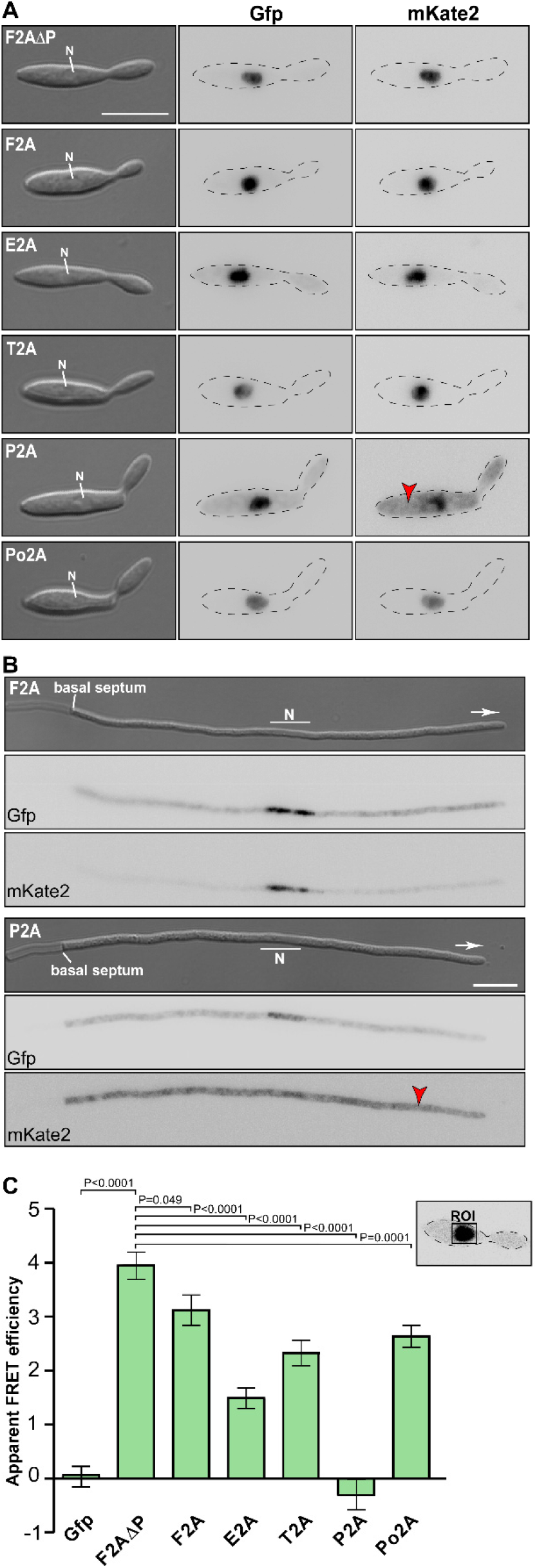
Separation efficiencies of 2A peptides analysed *in vivo*. Morphology and fluorescence microscopy of yeast (**A**) and hyphal (**B**) cells expressing reporter construct for analysis of 2A peptide separation efficiency (inverted fluorescence micrograph; N, nucleus; strong fluorescence signal in cytoplasm is indicated by red arrowhead; scale bar 10 μm; for hyphal cells: 6 h.p.i.; growth direction is indicated by arrow). (**C**) Apparent FRET efficiencies of tested 2A peptides determined within nucleus (ROI, region of interest; error bars, SEM; n = 3 independent experiments, for each experiment > 10 yeast cells were analysed; unpaired, two-tailed Mann-Whitney test).

Finally, we assessed the efficiency of the separation activity of the five different 2A peptides *in vivo* utilizing the read-out of fluorescence resonance energy transfer (FRET) between the reporters Gfp and mKate2 (Szymczak et al., 2004; characterisation of Gfp and mKate2: https://www.fpbase.org/compare/egfp,mkate2). The phenomenon of FRET can only occur if the donor fluorophore (Gfp^NLS^) is in very close proximity (below 10 nm) to the acceptor (mKate2^HA^). The further the two fluorescence proteins are separated from each other the less FRET is detectable. Thus, FRET experiments conducted in the nucleus reveal the proportion of reporter proteins that are not separated (Figure 3C), because the unseparated reporter fusion proteins mKate^HA^-2A-Gfp^NLS^ would accumulate in this compartment due to the presence of the NLS. Note that mKate^HA^ is able to enter the nucleus due to its small size even in the absence of an NLS (Figure 3B). However, a so called “bystander FRET” effect (unspecific FRET due to crowding of non-interacting donor and acceptor) is only expected at very high concentrations of proteins. Thus, a small amount of free mKate2 protein as observed in this case does not interfere with the measurement.

The experimental set-up revealed different low-level, but very stable FRET effects in the investigated 2A samples. The highest apparent FRET efficiency was observed in the nucleus of cells expressing the unseparated negative control F2AΔP, contrary to reduced FRET_app_ efficiencies within cells containing the assemblies with the different 2A peptides (Figure 3C). This underlines the sensitivity of the FRET measurements.

The experimental set-up was sensitive enough to detect the low F2A activity *in vivo* (Figure 3C). P2A shows FRET rates, which were comparable to a control strain only expressing Gfp, emphasizing a nearly 100% separation rate determined *in vivo*. The negative value of FRET_app_ is due to slight acquisition bleaching of Gfp. This is consistent with the results indicated by the Western blot analysis (Figure 2B) and the live cell fluorescence microscopy (Figure 3A-B). In essence, using a sophisticated *in vivo* strategy we were able to show that P2A exhibits the highest separation efficiency for *U. maydis*. Thus, we used P2A for further applications. When comparing different organisms, it is evident that 2A peptides exhibit a wide range of activities (Table 1). This underlines the importance of testing various 2A peptides regarding their separation efficiency, although P2A works best in most systems tested so far.

### Applying 2A Peptide Technology to Monitor Regulated Gene Expression Over Time

To illustrate the applicability, we designed a strategy for an efficient read-out for monitoring regulated gene expression. We aimed to quantify induction of a promoter over time using straight forward reporter enzyme activity.

To this end, we combined the *Photinus pyralis* Firefly luciferase (FLuc; L. Hüsemann, N. Heucken, and M. Zurbriggen, manuscript in preparation) with Rrm4-Gfp on a bi-cistronic mRNA using the P2A peptide sequence and the L2 linker (Figure 4A). A luciferase was successfully used before to determine P2A activity in *Aspergillus niger* (Schuetze and Meyer, 2017). The activity of Firefly luciferase can easily be detected by adding the substrate luciferin to the cells. As an example for regulated expression, we employed the promoter P_crg1_, which is active in the presence of arabinose and inactive in the sole presence of glucose (Figure 4A; Brachmann et al., 2001). The construct was integrated at the *ip^S^* locus by homologous recombination in the genetic background of AB33rrm4Δ (Loubradou et al., 2001; Materials and methods). Western blot experiments of hyphae growing for six hours under uninduced and induced conditions revealed that the luciferase as well as Rrm4-Gfp were expressed in arabinose-containing medium and, as expected, both were fully separated (Figure 4B). Analysing hyphal growth in glucose and arabinose revealed that only in glucose-containing medium hyphae grew in a bipolar mode; this aberrant growth form is characteristic for loss of Rrm4 (Figure 4C; Becht et al., 2006). In medium containing arabinose, however, the cells grew unipolarly as expected (Figure 4C). Studying dynamic subcellular localisation demonstrated that endosomal shuttling of Rrm4-Gfp was not influenced by the carbon source (arabinose or glucose) in the control strain expressing Rrm4-Gfp at the native locus (AB33P_rrm4_Rrm4-Gfp). However, no fluorescence signal of Rrm4-Gfp was detected in the presence of glucose (Figure 4D), indicating that the promoter is switched off.

**Figure 4.**
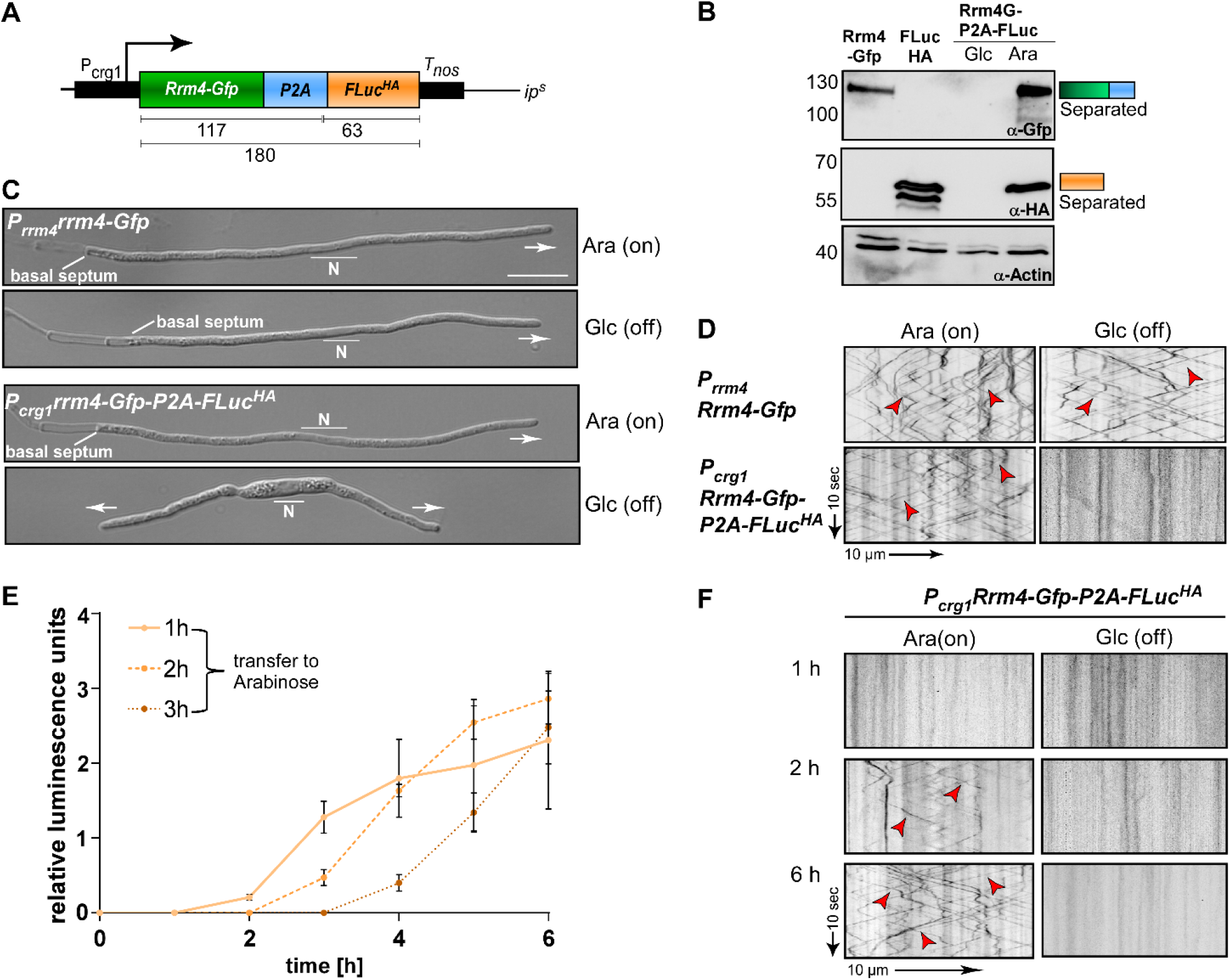
Following regulated gene expression over time using peptide P2A. (**A**) Schematic representation of a construct to analyse regulated gene expression (expected size of proteins in kDa given below). Arabinose inducible promoter (P_crg1_), Rrm4 fused to Gfp (green), L2 linker fused to P2A (blue), Firefly luciferase (FLuc) fused to HA tag (orange), transcriptional terminator (T_nos_). (**B**) Western blot analysis of AB33 derivates under induced and uninduced conditions (Glc: Glucose; Ara: Arabinose; antibodies are given at the bottom, size of marker proteins in kDa at the left). (**C**) Morphology of growing hypha expressing Rrm4 and FLuc separated by P2A in induced and uninduced conditions (6 h.p.i.; Glc: Glucose, Ara: Arabinose; growth direction is indicated by arrow; N, nucleus; scale bar 10 μm). (**D**) Kymographs of AB33 hyphae (6 h.p.i.) expressing Rrm4 and FLuc separated by P2A in induced and uninduced conditions (Glc: Glucose; Ara: Arabinose; arrow length on the left and bottom indicates time and distance) Bidirectional movement is indicated with red arrowheads. (**E**) Luciferase activity determination of strains expressing Rrm4-Gfp-L2-P2A-FLuc^HA^ shifted to hyphal growth at time point 0. After 1, 2 or 3 h cells were transferred to arabinose-containing medium (error bars, SEM; n = 3 independent experiments, relative luminescence units are given. (**F**) Kymographs of hypha carrying construct Rrm4-Gfp-P2A-FLuc after different time points of switching to arabinose containing medium (Glc: Glucose; Ara: Arabinose; arrow length on the left and bottom indicates time and distance; bidirectional movement is indicated with red arrowheads).

To analyse the induction of the P_crg1_ promoter in the presence of arabinose, hyphal growth was elicited in the presence of glucose. At different times the cells were transferred into arabinose-containing medium and FLuc activity as well as the fluorescence signal of Rrm4-Gfp were determined. Luciferase activity increased after shifting to arabinose-containing medium indicating activation of the P_crg1_ promoter (Figure 4E). Consistently, dynamic live cell imaging showed that endosomal shuttling of Rrm4-Gfp was detectable after two hours of growth in arabinose-containing medium (Figure 4F). After six hours the signal intensities were comparable to a strain expressing Rrm4-Gfp under the control of the native promoter. Thus, using 2A peptide technology allows simple and reliable quantification of gene expression. This can be used to study other aspects like mRNA stability, protein turnover or degradation, as well as to monitor the expression of certain mRNAs *in planta*.

### Synthetic Constitutive Co-expression of MEL Cluster Enzymes with the P2A Peptide

For the synthesis of biosurfactants genes for several biosynthetic enzymes need to be expressed simultaneously. This occurs in wild type strains, when the nitrogen source is limited. Here, we tested the applicability of the 2A peptide for synthetic activation of a secondary metabolite gene cluster in a biotechnological approach. This offers the clear advantage that synthesis can be uncoupled from nitrogen metabolism. *U. maydis* is a natural producer of the glycolipids mannosylerythritol lipid (MEL) and Ustilagic acid (UA). The biosynthetic enzymes for MEL production are encoded in a gene cluster activated upon nitrogen starvation (Figure 5A; Hewald et al., 2006). To achieve synthetic activation from a strong constitutively active promoter (P_oma_), we designed a tri-cistronic messenger RNA with three enzymes of the pathway. We used P2A combined with the L2 linker, and integrated the construct at the *ip^s^* locus of the glycolipid producing strain MB215 lacking biosynthesis of UA (MB215rua1Δ; Figure 5B). Genes for the mannosyltransferase Emt1 and cytoplasmic versions of the acyl transferases Mac1 and Mac2 (Freitag et al., 2014) were encoded on a single mRNA. For protein detection different tags were used (Figure 5B). Western blot experiments confirmed production of the three enzymes as separated proteins, indicating that a tri-cistronic construct is also functional (Figure 5C). Analysis of the glycolipid profile using thin-layer chromatography resulted in the production of MEL variants already detectable after 12 hours of growth (Figure 5D; see Materials and methods).

**Figure 5.**
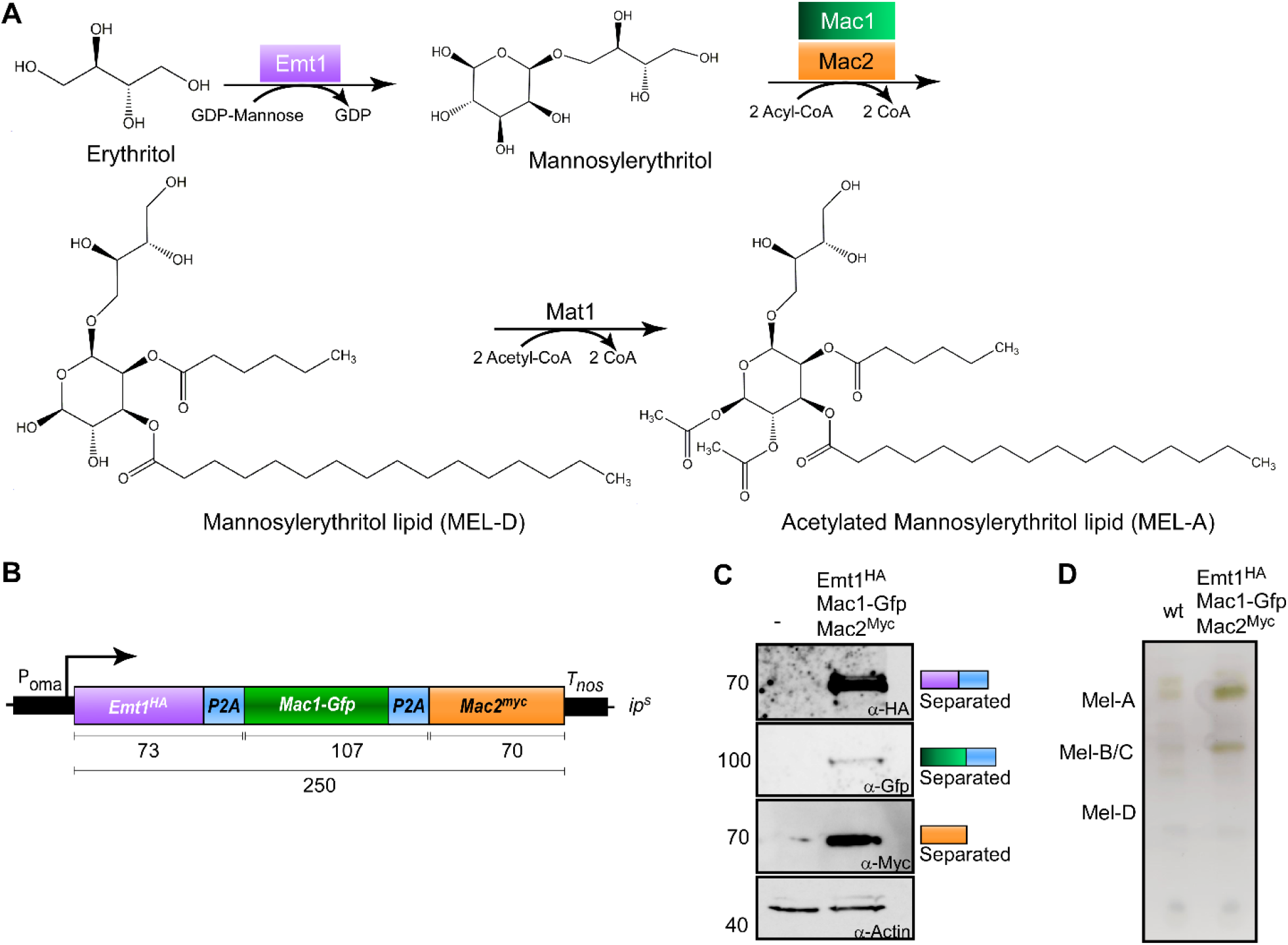
Usage of P2A enables production of biosurfactants. (**A**) Metabolic MEL synthesis pathway in *U. maydis*. GDP-mannose is added to an erythritol moiety by the mannosyl transferase Emt1 (purple) with consumption of GDP, resulting in the formation of mannosylerythritol. C_2_ and C_3_ positions are acylated with C_2_-C_18_ fatty acid chains. This step is catalysed by the acyl transferases Mac1 (green) and Mac2 (orange) and results in the formation of the deacetylated variant MEL-D. MEL-D is acetylated in positions C_4_ and C_6_ by the acetyl transferase Mat1, resulting in the formation of MEL-A (C_4_ and C_6_ acetylated), MEL-B (C_6_ acetylated, not shown) and MEL-C (C_4_ acetylated, not shown). (**B**) Schematic representation of tri-cistronic expression construct for MEL-D production: constitutively active promoter (P_oma_), ORF encoding Emt1 fused to HA tag (purple, Emt1^HA^), L2 fused to P2A (blue, P2A), Mac1 fused to Gfp (green, Mac1-Gfp), L2 fused to P2A (blue, P2A) Mac2 fused to a myc tag (orange, Mac2^myc^), transcriptional terminator (T_nos_). (**C**) Western blot analysis of protein synthesis via tri-cistronic mRNA (antibodies are given at the bottom, size of marker proteins in kDa at the left). (**D**) Thin layer chromatography of glycolipids from whole cells (progenitor strain MB215: wt).

The strategy to use 2A peptides to co-express multiple genes of biosynthetic pathways has successfully been used before. (i) Carotenoids were produced in *S. cerevisiae* (Beekwilder et al., 2014), (ii) β-lactam antibiotics or (iii) psychotropic mushroom alkaloids in *A. nidulans* (Unkles et al., 2014;Hoefgen et al., 2018}) as well as (iv) fungal toxins in *A. niger* (Schuetze and Meyer, 2017). This approach has now been expanded to the basidiomycete *U. maydis* underlining the broad applicability of polycistronic mRNAs in biotechnology.

### Conclusion

Here we present a straightforward strategy to analyse and quantitatively assess the functionality of 2A peptides *in vivo*. The analysis was conducted with five different versions but can easily be extended to other 2A peptides. The initial fluorescence readout of cytoplasmic red fluorescence is very simple and the FRET approach allows a sensitive and quantitative measurement of the separation activity. We successfully applied the best performing P2A peptide in basic and applied science demonstrating its efficient performance. With this proof-of-principle in hand, numerous new future applications like defined co-expression of subunits of protein complexes as well as efficient expression of heterologous biosynthetic gene clusters are conceivable.

## Materials and methods

### Plasmids, Strains and Growth Conditions

For cloning of plasmids, *E. coli* Top10 cells (Life Technologies, Carlsbad, CA, USA) were used. Transformation, cultivation and plasmid isolation were performed using standard techniques. *U. maydis* strains either derive from the lab strain AB33, in which the hyphal growth can be induced (Brachmann et al., 2001) or from the wild type strain MB215. AB33: Yeast-like cells were grown in complete medium (CM) supplemented with 1% glucose. Hyphal growth was induced by switching the nitrogen source by changing the media to nitrate minimal medium (NM) supplemented with 1% glucose or arabinose. MB215: MEL production cultures were incubated in unbaffled 300 ml flasks in 20 ml Verduyn-C mineral medium (0.28 M (5%) glucose, 0.23 M NH_4_NO_3_, 0.1 M MES pH 6.5, 3.6 mM KH_2_PO_4_, 0.8 mM MgSO_4_*7H_2_O, 51 μM EDTA, 37 μM FeCL_3_*6H_2_O, 16 μM H_3_BO_3_, 15.6 μM ZnSO_4_*7H_2_O, 6.7 μM MnCl_2_*2H_2_O, 2.3 μM CoCl_2_*6H_2_O, 1.9 μM Na_2_MoO_4_*2H_2_O, 0.6 μM KI) at 300 rpm and 28 °C. Cultures were inoculated to an OD_600_ of 0.1 and cultivated for up to 35 h.

Detailed growth conditions and cloning strategies for *U. maydis* are described elsewhere (Brachmann et al., 2004;Baumann et al., 2012;Terfrüchte et al., 2014;Beyer et al., 2015). All plasmids were sequenced to verify correctness. *U. maydis* strains were generated via homologous recombination within the *ip^S^* or the *upp3* locus (Loubradou et al., 2001) by transforming progenitor strains with linearised plasmids, except for MB215rua1Δ that was obtained with CRISPR-Cas technology (Schuster et al., 2016). Correctness of the strains was verified by counter selection (where possible), analytic PCR and Southern Blot analysis (Brachmann et al., 2004). A description of all plasmids and strains is summarised in Supplementary Table S1-S3. Sequences are available upon request.

### Microscopy, FRET and Image Analysis

For microscopy, yeast-like cells were grown for 12 h in complete medium. Microscopy was performed as described before (Baumann et al., 2012). The wide-field microscope Zeiss (Oberkochen, Germany) Axio Observer.Z1 provided with an Orca Flash4.0 camera (Hamamatsu, Japan) and objective lens Plan Apochromat (63x, NA 1.4) was used. Excitation of fluorescently-labelled proteins was carried out using a laser-based epifluorescence-microscopy. A VS-LMS4 Laser Merge-System (Visitron Systems, Puchheim, Germany) combines solid state lasers for the excitation of Gfp (488 nm/100 mW) and Rfp/mCherry (561 nm/150 mW). All modules of the microscope systems were controlled by the software package VisiView (Visitron). This was also used for image processing.

FRET-APB was measured using a Zeiss LSM780 laser-scanning microscope and a C-Apochromat 40x/1.20 korr M27 water objective (Carl Zeiss, Jena, Germany). GFP was excited with a 488 nm argon laser at an output power of 0.3% and emission of the fluorescence signal was detected between 490 to 552 nm using a 32 channel GaAsP detection unit. mKate2 was excited using a 561 nm DPSS laser at an output power of 5 % and emission detected between 588 to 686 nm. In total, a time series of 20 frames (256 times 256 pixels) at a pixel time of 3.15 μs/pixel was recorded with no line averaging. After the 5th frame, the nucleus and the surrounding area of yeast-like cells was bleached at 100 % laser power of the 561 nm laser, for 50-100 iterations. After the bleaching, 15 more frames were recorded. The “apparent FRET efficiency”, FRET_app_ was determined by comparing the fluorescence intensity in the bleached “region of interest” (ROI) of the donor fluorophore after bleaching of the acceptor fluorophore mKate2 according to the formula:

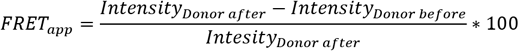

### Protein Extracts and Western Blot Analysis

*U. maydis* yeast-like cells or hyphae (6 h.p.i) were harvested by centrifugation (7,546 *g*, 10 minutes) and resuspended in 1 ml urea buffer (8M urea, 50 mM Tris/HCl pH8) to which protease inhibitors were freshly added (1 tablet of Complete protease inhibitor per 25 ml, Roche, Mannheim, Germany; 1 mM DTT; 0.1 M PMSF; 0.5 M benzamidine). After adding 200 μl of glass beads the cells were disrupted in 1.5 ml Eppendorf tubes with the Mixer Mill MM400 (Retsch, Haan, Germany) by agitating for 10 min at 30 Hz at 4 °C. For hyphae, the cells were agitated three times with cooling steps of 10 min in between. Protein concentrations were measured with the Bradford assay (Bio-Rad, Munich, Germany) and samples were adjusted to equal amounts. For Western Blot analysis, protein samples were supplemented with Laemmli buffer and heated to 95 °C for 6 min followed by centrifugation for 30 s at 16,200 g. Proteins were separated by 8, 10 or 12 % SDS-PAGE and transferred and immobilized in a n.itrocellulose membrane (GE Healthcare, Munich, Germany) by semi-dry blotting. Proteins were de gtected using α-Gfp, α-HA (both Roche, Freiburg, Germany), α-Myc (Sigma-Aldrich Chemie GmbH, Munich, Germany) and α-Actin (MP Biomedicals, Eschwege, Germany) antibodies. As secondary antibody an anti-mouse IgG HRP conjugate (Promega, Madison, WI, USA) was used. Detection was carried out by using Amersham™ ECL™ Prime (GE Healthcare, Munich, Germany). The images were taken according to the manual’s instructions with a luminescence image analyser, LAS4000 (GE Healthcare, Solingen, Germany).

### Luminescence measurements of firefly luciferase

To measure the luminescence of FLuc, 80 μl of cell suspension of hyphal growing cells was mixed with 20 μl of luciferin (20 mM Tricine, 2.67 mM MgSO_4_*7H2O, 0.1 mM EDTA*2H_2_O, 33.3 mM DTT, 524 μM ATP, 218 μM AcetylCoA, 131 μg/ml Luciferin, 5 mM NaOH, 264 μM MgCO_3_*5H_2_O) in a white Berthold 96-well plate (Nr: 23300/23302). The measurements lasted 20 minutes and were conducted using a BertholdTech Mithras luminescence reader (Berthold Technologies, Bad Wildbad, Germany) with the driver version 1.07.

### MEL extraction

MELs were sampled 12 h post inoculation. Therefore, 500 μl of whole cell culture broth were mixed with 500 μl of ethyl-acetate in 2 ml reaction tubes. MELs were extracted by shaking at 2,000 rpm for 15 min on a Vibrax VKA basic (IKA Werke GmbH & Co. KG, Staufen im Breisgau, Germany). Organic and aqueous phases were then separated by centrifugation at 21,100 *g* for 15 min. The organic phase was transferred into a fresh 1.5 ml reaction tube and evaporated at 70 °C for 1 h. Dried MELs were resolved in 15 μl methanol.

### MEL analysis by thin-layer chromatography

MEL production was analysed by TLC using a two-chamber system (modified from Hewald et al., 2006). Glycolipid extracts of up to 15 μl were applied evenly onto half TLC silica plates (20 x 10 cm, Merck KGaA, Darmstadt, Germany). After drying, plates were placed into a TLC chamber saturated with 100 ml buffer I (65:25:4 chloroform, methanol, H_2_O) for 5 min. Afterwards, plates were placed into a second TLC chamber saturated with 100 ml buffer II (9:1 chloroform, methanol) for 17 min. This step was repeated. For detection, dried TLC plates were sprayed with staining solution (50:1:0.5 glacial acetic acid, sulphuric acid, 4-methoxybenzaldehyde), dried again and incubated at 110 °C for 5 min.

## Supplementary Material

The Supplementary Material for this article can be found online at: https://www./xx.

## Data availability

All data generated or analysed during this study are included in the manuscript and/or the Supplementary Files.

## Author contributions

KM, MP, LH, NH, KS, MDZ and MF designed and planned the study. KM established the 2A peptides and analysed the promoter induction. MP optimized the production of the MELs. SWP and KM performed the FRET analysis. KM, KS and MF analysed the data. KM, KS, MDZ and MF designed and revised the manuscript. MF and KS directed the project.

## Funding

The work was funded by the Deutsche Forschungsgemeinschaft (DFG, German Research Foundation) Project FE448/9-1 to MF, Project-ID 267205415 – SFB 1208 to MDZ, SWP and MF as well as under Germany’s Excellence Strategy EXC-2048/1 – Project ID 39068111 to MDZ and MF). The scientific activities of the Bioeconomy Science Center were financially supported by the Ministry of Culture and Science within the framework of the NRW Strategieprojekt BioSC (No. 313/323-400-002 13 to MDZ and MF).

## Acknowledgements

We acknowledge lab members for discussion and comments on the manuscript. We are grateful to S. Kolar and B. Axler for excellent technical assistance, to Drs. M. Bölker and B. Sandrock for assistance with glycolipid analysis as well as providing the CRISPR plasmid for *rua1* deletion and P. Stoffels for support in strain generation. We thank Philipp Rink for preliminary work on 2A peptides for MEL synthesis.

## Notes

### Competing Interest Statement

The authors have declared no competing interest.

## References

Atkins, J.F., Wills, N.M., Loughran, G., Wu, C.Y., Parsawar, K., Ryan, M.D., Wang, C.H., and Nelson, C.C. (2007). A case for “StopGo”: reprogramming translation to augment codon meaning of GGN by promoting unconventional termination (Stop) after addition of glycine and then allowing continued translation (Go). RNA 13, 803–810.

Baumann, S., König, J., Koepke, J., and Feldbrügge, M. (2014). Endosomal transport of septin mRNA and protein indicates local translation on endosomes and is required for correct septin filamentation. EMBO Rep. 15, 94–102.

Baumann, S., Pohlmann, T., Jungbluth, M., Brachmann, A., and Feldbrügge, M. (2012). Kinesin-3 and dynein mediate microtubule-dependent co-transport of mRNPs and endosomes. J. Cell Sci. 125, 2740–2752.

Becht, P., König, J., and Feldbrügge, M. (2006). The RNA-binding protein Rrm4 is essential for polarity in *Ustilago maydis* and shuttles along microtubules. J. Cell Sci. 119, 4964–4973.

Becker, J., Hosseinpour Tehrani, H., Gauert, M., Mampel, J., Blank, L.M., and Wierckx, N. (2019). An *Ustilago maydis* chassis for itaconic acid production without by-products. Microb. Biotechnol.

Beekwilder, J., Van Rossum, H.M., Koopman, F., Sonntag, F., Buchhaupt, M., Schrader, J., Hall, R.D., Bosch, D., Pronk, J.T., Van Maris, A.J., and Daran, J.M. (2014). Polycistronic expression of a beta-carotene biosynthetic pathway in *Saccharomyces cerevisiae* coupled to beta-ionone production. J. Biotechnol. 192 Pt B, 383–392.

Béthune, J., Jansen, R.P., Feldbrügge, M., and Zarnack, K. (2019). Membrane-associated RNA-binding proteins orchestrate organelle-coupled translation Trends Cell Biol. 29, 178–188.

Beyer, H.M., Gonschorek, P., Samodelov, S.L., Meier, M., Weber, W., and Zurbriggen, M.D. (2015). AQUA cloning: a versatile and simple enzyme-free cloning approach. PLoS One 10, e0137652.

Brachmann, A., König, J., Julius, C., and Feldbrügge, M. (2004). A reverse genetic approach for generating gene replacement mutants in *Ustilago maydis*. Mol. Gen. Genom. 272, 216–226.

Brachmann, A., Weinzierl, G., Kämper, J., and Kahmann, R. (2001). Identification of genes in the bW/bE regulatory cascade in *Ustilago maydis*. Mol. Microbiol. 42, 1047–1063.

Brefort, T., Doehlemann, G., Mendoza-Mendoza, A., Reissmann, S., Djamei, A., and Kahmann, R. (2009). *Ustilago maydis* as a Pathogen. Annu. Rev. Phytopathol. 47, 423–445.

Chng, J., Wang, T., Nian, R., Lau, A., Hoi, K.M., Ho, S.C., Gagnon, P., Bi, X., and Yang, Y. (2015). Cleavage efficient 2A peptides for high level monoclonal antibody expression in CHO cells. MAbs 7, 403–412.

Cioni, J.M., Lin, J.Q., Holtermann, A.V., Koppers, M., Jakobs, M.a.H., Azizi, A., Turner-Bridger, B., Shigeoka, T., Franze, K., Harris, W.A., and Holt, C.E. (2019). Late endosomes act as mRNA translation platforms and sustain mitochondria in axons. Cell 176, 56–72 e15.

Daniels, R.W., Rossano, A.J., Macleod, G.T., and Ganetzky, B. (2014). Expression of multiple transgenes from a single construct using viral 2A peptides in *Drosophila*. PLoS One 9, e100637.

De Felipe, P., Luke, G.A., Hughes, L.E., Gani, D., Halpin, C., and Ryan, M.D. (2006). E unum pluribus: multiple proteins from a self-processing polyprotein. Trends Biotechnol. 24, 68–75.

De Felipe, P., and Ryan, M.D. (2004). Targeting of proteins derived from self-processing polyproteins containing multiple signal sequences. Traffic 5, 616–626.

Donnelly, M.L., Gani, D., Flint, M., Monaghan, S., and Ryan, M.D. (1997). The cleavage activities of aphthovirus and cardiovirus 2A proteins. J. Gen. Virol. 8 (Pt 1), 13–21.

Feldbrügge, M., Kellner, R., and Schipper, K. (2013). The biotechnological use and potential of plant pathogenic smut fungi. Appl. Microbiol. Biotechnol. 97, 3253–3265.

Freitag, J., Ast, J., Linne, U., Stehlik, T., Martorana, D., Bölker, M., and Sandrock, B. (2014). Peroxisomes contribute to biosynthesis of extracellular glycolipids in fungi. Mol. Microbiol. 93, 24–36.

Gao, S.Y., Jack, M.M., and O’neill, C. (2012). Towards optimising the production of and expression from polycistronic vectors in embryonic stem cells. PLoS One 7, e48668.

Geier, M., Fauland, P., Vogl, T., and Glieder, A. (2015). Compact multi-enzyme pathways in P. pastoris. Chem. Commun. 51, 1643–1646.

Geiser, E., Reindl, M., Blank, L.M., Feldbrügge, M., Wierckx, N., and Schipper, K. (2016). Activating intrinsic carbohydrate-active enzymes of the smut fungus *Ustilago maydis* for the degradation of plant cell call components. Appl. Environ. Microbiol. 82, 5174–5185.

Gibson, D.G., Young, L., Chuang, R.Y., Venter, J.C., Hutchison, C.A., 3rd, and Smith, H.O. (2009). Enzymatic assembly of DNA molecules up to several hundred kilobases. Nat. Methods 6, 343–345.

Ha, S.H., Liang, Y.S., Jung, H., Ahn, M.J., Suh, S.C., Kweon, S.J., Kim, D.H., Kim, Y.M., and Kim, J.K. (2010). Application of two bicistronic systems involving 2A and IRES sequences to the biosynthesis of carotenoids in rice endosperm. Plant Biotechnol. J. 8, 928–938.

Halpin, C., Cooke, S.E., Barakate, A., El Amrani, A., and Ryan, M.D. (1999). Self-processing 2A-polyproteins--a system for co-ordinate expression of multiple proteins in transgenic plants. Plant J. 17, 453–459.

Hewald, S., Linne, U., Scherer, M., Marahiel, M.A., Kämper, J., and Bölker, M. (2006). Identification of a gene cluster for biosynthesis of mannosylerythritol lipids in the basidiomycetous fungus *Ustilago maydis*. Appl. Environ. Microbiol. 72, 5469–5477.

Hoefgen, S., Lin, J., Fricke, J., Stroe, M.C., Mattern, D.J., Kufs, J.E., Hortschansky, P., Brakhage, A.A., Hoffmeister, D., and Valiante, V. (2018). Facile assembly and fluorescence-based screening method for heterologous expression of biosynthetic pathways in fungi. Metab. Eng. 48, 44–51.

Holst, J., Vignali, K.M., Burton, A.R., and Vignali, D.A. (2006). Rapid analysis of T-cell selection *in vivo* using T cell-receptor retrogenic mice. Nat. Methods 3, 191–197.

Kahmann, R., and Kämper, J. (2004). *Ustilago maydis*: how its biology relates to pathogenic development. New Phytol. 164, 31–42.

Keene, J.D. (2007). RNA regulons: coordination of post-transcriptional events. Nat. Rev. Genet. 8, 533–543.

Kim, J.H., Lee, S.R., Li, L.H., Park, H.J., Park, J.H., Lee, K.Y., Kim, M.K., Shin, B.A., and Choi, S.Y. (2011). High cleavage efficiency of a 2A peptide derived from porcine teschovirus-1 in human cell lines, zebrafish and mice. PLoS One 6, e18556.

König, J., Baumann, S., Koepke, J., Pohlmann, T., Zarnack, K., and Feldbrügge, M. (2009). The fungal RNA-binding protein Rrm4 mediates long-distance transport of *ubi1* and *rho3* mRNAs. EMBO J. 28, 1855–1866.

Liao, Y.C., Fernandopulle, M.S., Wang, G., Choi, H., Hao, L., Drerup, C.M., Patel, R., Qamar, S., Nixon-Abell, J., Shen, Y., Meadows, W., Vendruscolo, M., Knowles, T.P.J., Nelson, M., Czekalska, M.A., Musteikyte, G., Gachechiladze, M.A., Stephens, C.A., Pasolli, H.A., Forrest, L.R., St George-Hyslop, P., Lippincott-Schwartz, J., and Ward, M.E. (2019). RNA granules hitchhike on lysosomes for long-distance transport, using annexin A11 as a molecular tether. Cell 179, 147–164 e120.

Liu, Z., Chen, O., Wall, J.B.J., Zheng, M., Zhou, Y., Wang, L., Ruth Vaseghi, H., Qian, L., and Liu, J. (2017). Systematic comparison of 2A peptides for cloning multi-genes in a polycistronic vector. Sci. Rep. 7, 2193.

Loubradou, G., Brachmann, A., Feldbrügge, M., and Kahmann, R. (2001). A homologue of the transcriptional repressor Ssn6p antagonizes cAMP signalling in *Ustilago maydis*. Mol. Microbiol. 40, 719–730.

Müller, M.J., Stachurski, S., Stoffels, P., Schipper, P., Feldbrügge, M., and Büchs, J. (2018). Online evaluation of the metabolic activity of *Ustilago maydis* on (poly)galacturonic acid.. J. Biol. Eng. 34, e1–17.

Olgeiser, L., Haag, C., Boerner, S., Ule, J., Busch, A., Koepke, J., König, J., Feldbrügge, M., and Zarnack, K. (2019). The key protein of endosomal mRNP transport Rrm4 binds translational landmark sites of cargo mRNAs. EMBO Rep. 20, e46588.

Pohlmann, T., Baumann, S., Haag, C., Albrecht, M., and Feldbrügge, M. (2015). A FYVE zinc finger domain protein specifically links mRNA transport to endosome trafficking. Elife 4, e06041.

Provost, E., Rhee, J., and Leach, S.D. (2007). Viral 2A peptides allow expression of multiple proteins from a single ORF in transgenic zebrafish embryos. Genesis 45, 625–629.

Ryan, M.D., King, A.M., and Thomas, G.P. (1991). Cleavage of foot-and-mouth disease virus polyprotein is mediated by residues located within a 19 amino acid sequence. J. Gen. Virol. 72 (Pt 11), 2727–2732.

Sarkari, P., Reindl, M., Stock, J., Müller, O., Kahmann, R., Feldbrügge, M., and Schipper, K. (2014). Improved expression of single-chain antibodies in *Ustilago maydis*. J. Biotechnol. 191, 165–175.

Schuetze, T., and Meyer, V. (2017). Polycistronic gene expression in *Aspergillus niger*. Microb. Cell Fact. 16, 162.

Schuster, M., Schweizer, G., Reissmann, S., and Kahmann, R. (2016). Genome editing in *Ustilago maydis* using the CRISPR-Cas system. Fungal Genet. Biol. 89, 3–9.

Sharma, P., Yan, F., Doronina, V.A., Escuin-Ordinas, H., Ryan, M.D., and Brown, J.D. (2012). 2A peptides provide distinct solutions to driving stop-carry on translational recoding. Nucleic Acids Res. 40, 3143–3151.

Shcherbo, D., Merzlyak, E.M., Chepurnykh, T.V., Fradkov, A.F., Ermakova, G.V., Solovieva, E.A., Lukyanov, K.A., Bogdanova, E.A., Zaraisky, A.G., Lukyanov, S., and Chudakov, D.M. (2007). Bright far-red fluorescent protein for whole-body imaging. Nat. Methods 4, 741–746.

Shcherbo, D., Murphy, C.S., Ermakova, G.V., Solovieva, E.A., Chepurnykh, T.V., Shcheglov, A.S., Verkhusha, V.V., Pletnev, V.Z., Hazelwood, K.L., Roche, P.M., Lukyanov, S., Zaraisky, A.G., Davidson, M.W., and Chudakov, D.M. (2009). Far-red fluorescent tags for protein imaging in living tissues. Biochem. J. 418, 567–574.

Souza-Moreira, T.M., Navarrete, C., Chen, X., Zanelli, C.F., Valentini, S.R., Furlan, M., Nielsen, J., and Krivoruchko, A. (2018). Screening of 2A peptides for polycistronic gene expression in yeast. FEMS Yeast Res. 18.

Steinberg, G., and Perez-Martin, J. (2008). *Ustilago maydis*, a new fungal model system for cell biology. Trends Cell Biol. 18, 61–67.

Stoffels, P., Müller, M.J., Stachurski, S., Terfrüchte, M., Schroder, S., Ihling, N., Wierckx, N., Feldbrügge, M., Schipper, K., and Büchs, J. (2020). Complementing the intrinsic repertoire of *Ustilago maydis* for degradation of the pectin backbone polygalacturonic acid. J. Biotechnol. 307, 148–163.

Subramanian, V., Schuster, L.A., Moore, K.T., Taylor, L.E., 2nd, Baker, J.O., Vander Wall, T.A., Linger, J.G., Himmel, M.E., and Decker, S.R. (2017). A versatile 2A peptide-based bicistronic protein expressing platform for the industrial cellulase producing fungus, *Trichoderma reesei*. Biotechnol. Biofuels 10, 34.

Szymczak, A.L., Workman, C.J., Wang, Y., Vignali, K.M., Dilioglou, S., Vanin, E.F., and Vignali, D.A. (2004). Correction of multi-gene deficiency in vivo using a single ‘self-cleaving’ 2A peptide-based retroviral vector. Nat. Biotechnol. 22, 589–594.

Teichmann, B., Liu, L., Schink, K.O., and Bolker, M. (2010). Activation of the ustilagic acid biosynthesis gene cluster in *Ustilago maydis* by the C2H2 zinc finger transcription factor Rua1. Appl. Environ. Microbiol. 76, 2633–2640.

Terfrüchte, M., Joehnk, B., Fajardo-Somera, R., Braus, G., Riquelme, M., Schipper, K., and Feldbrügge, M. (2014). Establishing a versatile Golden Gate cloning system for genetic engineering in fungi. Fungal Genet. Biol. 62, 1–10.

Unkles, S.E., Valiante, V., Mattern, D.J., and Brakhage, A.A. (2014). Synthetic biology tools for bioprospecting of natural products in eukaryotes. Chem. Biol. 21, 502–508.

Wang, Y., Wang, F., Wang, R., Zhao, P., and Xia, Q. (2015). 2A self-cleaving peptide-based multi-gene expression system in the silkworm *Bombyx mori*. Sci. Rep. 5, 16273.

Zarnack, K., Maurer, S., Kaffarnik, F., Ladendorf, O., Brachmann, A., Kämper, J., and Feldbrügge, M. (2006). Tetracycline-regulated gene expression in the pathogen *Ustilago maydis*. Fungal Genet. Biol. 43, 727–738.

Zhou, L., Obhof, T., Schneider, K., Feldbrügge, M., Nienhaus, G.U., and Kämper, J. (2018). Cytoplasmic transport machinery of the SPF27 homologue Num1 in *Ustilago maydis*. Sci. Rep. 8, 3611.

