## Supplementary Material for "Polycistronic gene expression in the model micro-organism *Ustilago maydis*"

Including Supplementary Figure S1 as well as Supplementary Table S1-S3

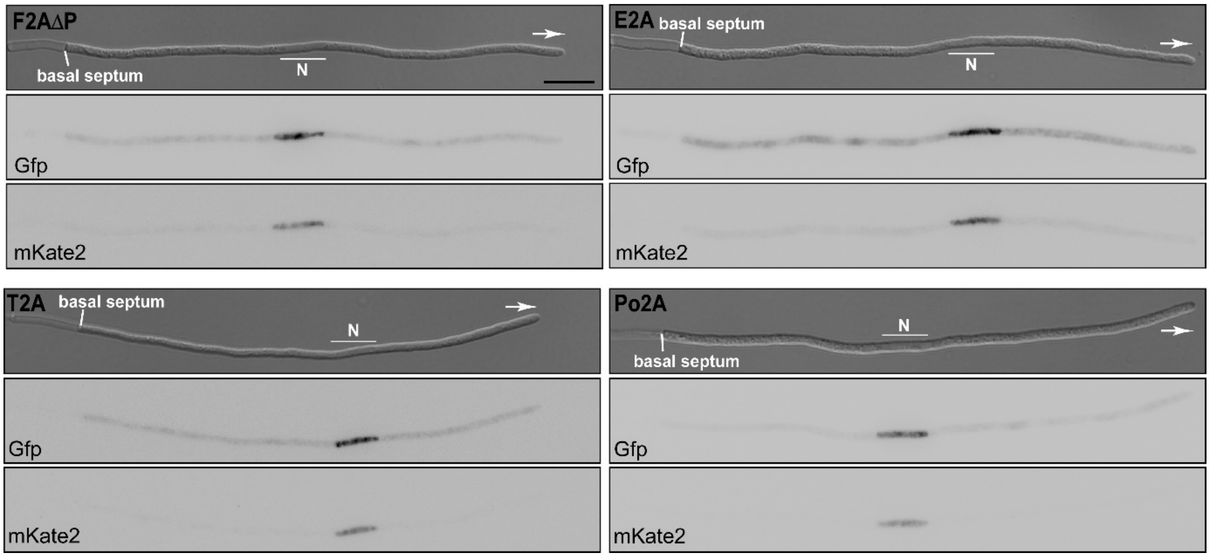


**Supplementary Figure S1. Separation efficiency of 2A peptides during hyphal growth *in vivo.***

Hyphal cells (6 h.p.i.) expressing reporter construct mKate2$^{HA}$-L2-2A-Gfp$^{NLS}$ (inverted fluorescence micrographs; N, nucleus; scale bar 10 µm; growth direction is indicated by arrow).

**Supplementary Table S1: *U. maydis* strains used in this study**; UMa, internal reference number

| Strain | Relevant genotype | Short description | Uma | Reference |
|---|---|---|---|---|
| **AB33** | a2 P*nar:bWebE1* | expression of active b heterodimer which is under the control of $P_{nar1}$ promoter; hyphal growth can be induced by switching the nitrogen source | 133 | (Brachmann, 2001) |
| **AB33*rrm4*Δ** | *rrm4*Δ | carries a deletion of *rrm4* | 273 | (Becht et al., 2006) |
| **AB33Rrm4-Gfp** | *rrm4-Gfp* | expresses Rrm4 C-terminally fused to Gfp | 274 | (Becht et al., 2006) |
| **AB33Rrm4-mCherry** | *rrm4-mCherry* | expresses Rrm4 C-terminally fused to mCherry | 830 | (Baumann et al., 2014) |
| **AB33Rrm4-TagRfp** | *rrm4-TagRfp* | expresses Rrm4 C-terminally fused to TagRfp | 1317 | (Müntjes, 2015) |
| **AB33Rrm4-mKate2** | *rrm4-mKate2* | expresses Rrm4 C-terminally fused to mKate2 | 1985 | This study |
| **AB33*upp3*Δ::$P_{otef}$-mKate2-HA-GSG-F2A-Gfp-NLS** | *mKate2-GSG-F2A-Gfp-NLS* | co-expresses mKate2 fused to a HA tag and Gfp fused to an NLS partial separated during translation by F2A | 2495 | This study |
| **AB33*upp3*Δ::$P_{otef}$-mKate2-HA-F2AΔP-Gfp-NLS** | *mKate2-F2AΔP-Gfp-NLS* | expresses a fusion protein of mKate2 fused to a HA tag and Gfp fused to an NLS; no separation during translation because of deletion of last proline of F2A | 2520 | This study |
| **MB215 rua1*Δ*** | *rua1Δ (CRISPR)* | carries a deletion of *rua1* | 2598 | (Peter Stoffels, unpublished) |
| **AB33*upp3*Δ::$P_{otef}$-Firefly-HA** | *firefly-HA* | expresses Firefly luciferase fused to an HA tag | 2644 | (L. Hüsemann et al., in preparation) |
| **MB215*rua1Δ*/$P_{oma}$-*emt*1-HA-L2-P2A-Gfp-*mac*1-L2-P2A-*mac*2-3xmyc** | *emt1-HA-L2-P2A-gfp-mac1-L2-P2A-mac2-3xmyc* | co-expresses Emt1 fused to an HA tag, Mac1 N-terminally fused to Gfp and Mac2 fused to a triple myc tag; all proteins are separation by P2A fused N-terminally to L2 linker during translation | 3084 | This study |
| **AB33*upp3*Δ::$P_{otef}$-mKate2-HA-L1-F2A-Gfp-NLS** | *mKate2-L1-F2A-Gfp-NLS* | co-expresses mKate2 fused to a HA tag and Gfp fused to an NLS; partial separated during translation by F2A fused N-terminally to L1 linker | 3116 | This study |
| **AB33*upp3*Δ::$P_{otef}$-mKate2-HA-L2-F2AΔP-Gfp-NLS** | *mKate2-L2-F2AΔp-Gfp-NLS* | expresses fusion of mKate2 fused to a HA tag and Gfp fused to an NLS; no separation during translation because of deletion of last proline of F2A | 3119 | This study |
| **AB33*upp3*Δ::$P_{otef}$-mKate2-HA-L2-T2A-Gfp-NLS** | *mKate2-L2-T2A-Gfp-NLS* | co-expresses mKate2 fused to a HA tag and Gfp fused to an NLS partial separated during translation by T2A fused N-terminally to L2 linker | 3120 | This study |
| **AB33*upp3*Δ::$P_{otef}$-mKAte2-HA-L1-F2AΔP-Gfp-NLS** | *mKate2-L1-F2AΔlastp-Gfp-NLS* | expresses fusion protein of mKate2 fused to a HA tag and Gfp fused to an NLS; no separation during translation because of deletion of last proline of F2A | 3122 | This study |

| | | | | |
|---|---|---|---|---|
| **AB33*upp3*Δ::P$_{otef}$-mKate2-HA-L2-P2A-Gfp-NLS** | *mKate2-L2-P2A-Gfp-NLS* | co-expresses mKate2 fused to a HA tag and Gfp fused to an NLS separated during translation by P2A fused N-terminally to L2 linker | 3123 | This study |
| **AB33*upp3*Δ::P$_{otef}$-mKate2-HA-L2-F2A-eGFP-NLS** | *mKate2-L2-F2A-Gfp-NLS* | co-expresses mKate2 fused to a HA tag and Gfp fused to an NLS partial separated during translation by F2A fused to L2 linker | 3144 | This study |
| **AB33*upp3*Δ::P$_{otef}$-mKate2-HA-L2-Po2A-eGFP-NLS** | *mKate2-L2-Po2A-Gfp-NLS* | co-expresses mKate2 fused to a HA tag and Gfp fused to an NLS partial separated during translation by Po2A fused to L2 linker | 3145 | This study |
| **AB33*rrm4*Δ/Rrm4-Gfp-L2-P2A-Firefly-HA** | *rrm4-Gfp-L2-P2A-firefly-HA* | co-expresses Rrm4 fused C-terminally to Gfp and Firefly luciferase fused to HA tag; separation during translation through P2A N-terminally fused to L2 | 3212 | This study |

**Supplementary Table S2: *U. maydis* strains generated in this study**;
UMa and pUMa, internal reference number

| Strain | UMa | Relevant genotype | Transformed plasmid | Locus | Progenitor strain |
|---|---|---|---|---|---|
| **AB33Rrm4-mKate2** | 1985 | *Rrm4-mKate2* | $P_{rrm4}$-*rrm4*-mKate2 (pUMa2985) | *rrm4* | AB33*rrm4*Δ |
| **AB33*upp3*Δ::$P_{otef}$-mKate2-HA-F2A-Gfp-NLS** | 2495 | *mKate2-F2A-Gfp-NLS* | p*upp3*Δ::$P_{otef}$-mKate2-HA-F2A-Gfp-NLS (pUMa3407) | *upp3* | AB33*upp3*Δ |
| **AB33*upp3*Δ::$P_{otef}$-mKate2-HA-F2AΔP-Gfp-NLS** | 2520 | *mKate2-F2AΔP-Gfp-NLS* | p*upp3*Δ::$P_{otef}$-mKate2-HA-F2AΔP-Gfp-NLS (pUMa3435) | *upp3* | AB33*upp3*Δ |
| **MB215 *rua1Δ*/$P_{oma}$-*emt*1-HA-L2-P2A-Gfp-*mac*1-L2-P2A-*mac*2-3xmyc** | 3084 | *emt1-HA-L2-P2A-gfp-mac1-L2-P2A-mac2-3xmyc* | pip$^R$::$P_{oma}$-*emt*1-HA-L2-P2A-Gfp-*mac*1-L2-P2A-*mac*2-3xmyc (pUMa4131) | *ip$^S$* | MB215*rua1Δ* |
| **AB33*upp3*Δ::$P_{otef}$-mKate2-HA-L1-F2A-Gfp-NLS** | 3116 | *mKate2-L1-F2A-Gfp-NLS* | p*upp3*Δ::$P_{otef}$-mKate2-HA-L1-F2A-Gfp-NLS (pUMa4261) | *upp3* | AB33*upp3*Δ |
| **AB33*upp3*Δ::$P_{otef}$-mKate2-HA-L2-F2AΔp-Gfp-NLS** | 3119 | *mKate2-L2-F2AΔp-Gfp-NLS* | p*upp3*Δ::$P_{otef}$-mKate2-HA-L2-F2AΔP-Gfp-NLS (pUMa4197) | *upp3* | AB33*upp3*Δ |
| **AB33*upp3*Δ::$P_{otef}$-mKate2-HA-L2-T2A-Gfp-NLS** | 3120 | *mKate2-L2-T2A-Gfp-NLS* | p*upp3*Δ::$P_{otef}$-mKate2-HA-L2-T2A-Gfp-NLS (pUMa4202) | *upp3* | AB33*upp3*Δ |
| **AB33*upp3*Δ::$P_{otef}$-mKAte2-HA-L1-F2AΔp-Gfp-NLS** | 3122 | *mKate2-L1-F2AΔlastp-Gfp-NLS* | p*upp3*Δ::$P_{otef}$-mKate2-HA-L1-F2AΔP-Gfp-NLS (pUMa4273) | *upp3* | AB33*upp3*Δ |
| **AB33*upp3*Δ::$P_{otef}$-mKate2-HA-L2-P2A-Gfp-NLS** | 3123 | *mKate2-L2-P2A-Gfp-NLS* | p*upp3*Δ::$P_{otef}$-mKate2-HA-L2-P2A-Gfp-NLS (pUMa4199) | *upp3* | AB33*upp3*Δ |
| **AB33*upp3*Δ::$P_{otef}$-mKate2-HA-L2-F2A-eGFP-NLS** | 3144 | *mKate2-L2-F2A-Gfp-NLS* | p*upp3*Δ::$P_{otef}$-mKate2-HA-L2-F2A-Gfp-NLS (pUMa4198) | *upp3* | AB33*upp3*Δ |
| **AB33*upp3*Δ::$P_{otef}$-mKate2-HA-L2-Po2A-eGFP-NLS** | 3145 | *mKate2-L2-Po2A-Gfp-NLS* | p*upp3*Δ::$P_{otef}$-mKate2-HA-L2-Po2A-Gfp-NLS (pUMa4204) | *upp3* | AB33*upp3*Δ |
| **AB33*rrm4*Δ/Rrm4-Gfp-L2-P2A-Firefly-HA** | 3212 | *rrm4-Gfp-L2-P2A-firefly-HA* | p*ip*$^R$:$P_{crg}$-*rrm4*-Gfp-L2-P2A-*firefly*-HA (pUMa4565) | *ip$^S$* | AB33*rrm4*Δ |

**Appendix Table S3: Plasmids generated in this study**; pUMa, internal reference number

| Plasmid | pUMa | Resistance cassette | Short description |
|---|---|---|---|
| **$P_{rrm4}$-*rrm4*-mKate2** | 2985 | NatR (SfiI-insert of pMF5-1n) | Vector for the expression of Rrm4 C-terminally fused to mKate2. The mKate2 cassette contains the Tnos terminator and the Nat resistance. The entire coding sequence for the fusion protein is flanked by an 830 bp upstream region and a 1.9 kb downstream region for homologous recombination. The plasmid is a derivate of pRrm4G-NatR (Becht et al., 2006), with Gfp was exchanged to mKate2. |
| **p*upp3*Δ::$P_{otef}$-mKate2-HA-F2A-Gfp-NLS** | 3407 | NatR | Vector contains a fusion of mKate2-HA and Gfp-NLS. mKate2 with F2A in reverse overhang was amplified with ofw: ATGGTGTCGGAGCTCATC and orev: TTTGAGGAGATCGAAGTTGAGCAGCTGTTTCACGGGG GCGTAGTCGGGCACGTCGTAAGGGTAGAGCGGACCCTGCATATGGC GGTGACCG. Gfp with F2A in forward overhang was amplified with ofw: TGAAACAGCTGCTCAACTTCGATCTCCTCAAACTGGCCGG CGACGTGGAATCAAATCCTGGACCTATGGTGAGCAAGGGC and orev: CTTGTACAGCTCGTCCATG. In between a F2A peptide ensures expression of two proteins from one open reading frame Expression is driven by the constitutive $P_{otef}$ and terminated by $T_{nos}$. Flanking regions of 800 bp upstream and 700 bp downstream of the entire construct ensure homologous recombination at *upp3* locus. Plasmid was generated using AQUA cloning. |
| **p*upp3*Δ::$P_{otef}$-mKate2-HA-F2AΔP-Gfp-NLS** | 3435 | NatR | Same as pUMa3407. mKate2 with F2A in reverse overhang was amplified with ofw: ATGGTGTCGGAGCTCATC and orev:TTTGAGGAGATCG AAGTTGAGCAGCTGTTTCACGGGGGCGTAGTCGGGCACGTCGTAAG GGTAGAGCGGACCCTGCATATGGCGGTGACCG. Gfp with F2A in forward overhang was amplified with ofw:TGAAACAGCTGCTCA ACTTCGATCTCCTCAAACTGGCCGGCGACGTGGAATCAAATCCTGGA ATGGTGAGCAAGGGC and orev: CTTGTACAGCTCGTCCATG. In between a F2A is inserted in which the last proline is deleted. Plasmid was generated using AQUA cloning. |
| **pip$^{R}$::$P_{oma}$-*emt*1-HA-L2-P2A-Gfp-*mac*1-L2-P2A-*mac*2-3xmyc** | 4131 | CbxR (integration of *ip*$^{R}$ gene into *ip*$^{S}$ locus; ip: Iron-sulfur subunit of the suc-cinate dehydrogenase) | Vector for the expression of Emt1, Mac1 and Mac2 with respective tags, linked by the L2 linker and P2A in a single open reading frame. Emt1-HA with L2-P2A in reverse overhang was amplified with ofw:TACCTTACT CTATCAGGATCCCCGCATGAAGGTTGCCC TCCTTGC and orev: GCTCCTCGCCCTTGCTCACCATGGGACCGGGGTTCTCCTCGACGTCA CCG. *gfp-mac1* with P2A in forward overhang and L2 in reverse overhang was amplified with ofw:CGGTGACGTCGAGGAGACC CCGGTCCCATGGTGAGCAAGGGCGAGGAGC and orev:GCGGG AAGCCGTGGCTAAGCTTAAACACAGGAGCGGTCGCATCGACCG. Mac2 with L2-P2A in forward overhang and 3xmyc and Tnos in reverse overhang was amplified with ofw:ATGCGACCGCTCCTGTG TTTAAG CTTAGCCACGGCTTCCCGC and orev: GCAAAAGCGAAACAGCGG CGCGACCCTAGAGGTCCTCTTCCGAGATGAGCT TCTGCT CGGA CCACCGGCCGAGAGGTCCTCTTCCGAGATGAGCTTCTGCTCCGAGCC GGCACCGGCGAGGTCCTCTTCCGAGATGAGCTTCTGCTCCTCGGGA GCGTCAACGGGGACTG. The P2A peptide ensures translation of three proteins from a single open reading frame. Expression is driven by the synthetic constitutive promoter $P_{oma}$ and terminated by $T_{nos}$. The expression cassette contains the ip*R* gene, which is cleaved to ensure homologous recombination prior to transformation. |

| | | | |
|---|---|---|---|
| **p*upp3*Δ::P$_{otef}$-mKate2-HA-L1-F2A-Gfp-NLS** | 4261 | NatR | Vector contains a fusion of mKate2-HA and Gfp-NLS. mKate2 was amplified with ofw:GGTCTCGCCTGCCCTGCAGGCTAGAACT AGTG and orev:GGTCTCCAGGCCGGCCGGCGTAGTCGGGCA CGTCGTAAGG. Gfp was amplified with ofw:GGTCTCCGCCTGC GAGAGAGAGAGCTCAGCCCATGGTGAGCAAGGGCGAGG and orev: GGTCTCGCTGCGGCGCGCCGGCCGCTAGATC. In between a F2A peptide N-terminally fused to an GSG linker (L1) ensures expression of two proteins from one open reading frame. GSG-F2A was amplified with ofw:GGCCGGCCTGGCTCGGGCCCCGTGAAACAGCTGCT CAAC and orev:GGCCGGCCTCACGGCTTCCCGCCGGCGGTGG CGGCGCAGGATGATGGCACGCTGGTCGCCATGACCGTCATGGCCTTC . Expression is driven by the constitutive P$_{otef}$ and terminated by T$_{nos}$. Flanking regions of 800 bp upstream and 700 bp downstream of the entire construct ensure homologous recombination at *upp3* locus. |
| **p*upp3*Δ::P$_{otef}$-mKate2-HA-L2-F2AΔP-Gfp-NLS** | 4197 | NatR | Same as pUMa3261. Within F2A the last proline is deleted and it is N-terminally fused to KLSHGFPPAVAAQDDGTLV (L2). F2AΔP was amplified with ofw:GGCCGGCCTCACGGCTTCCCGCCGGCGGTGG CGGCGCAGGATGATGGCACGCTGGTCCCCGTGAAACAGCTGCTCAA C and orev: GCTGAGCTCCAGGATTTGATTCCACG. |
| **p*upp3*Δ::P$_{otef}$-mKate2-HA-L2-T2A-Gfp-NLS** | 4202 | NatR | Same as pUMa4197. T2A was amplified with ofw:GGCCGGCCTCACGG CTTCCCGCCGGCGGTGGCGGCGCAGGATGATGGCACGCTGGTCCGT GGTCCTCGTCCCCAGAAC and orev:GCTGAGCGGGACCGGGGT TCTCCTCGAC. |
| **p*upp3*Δ::P$_{otef}$-mKate2-HA-L1-F2AΔP-Gfp-NLS** | 4273 | NatR | Same as pUMa4261. F2ADP was amplified with ofw:GGCCGGCCTGGC TCGGGCCGCCACAAGTTCCCCACCAAC and orev:GCTGAGCGGG ACCGGGGTTCGACTCGAC. |
| **p*upp3*Δ::P$_{otef}$-mKate2-HA-L2-P2A-Gfp-NLS** | 4199 | NatR | Same as pUMa4197. P2A was amplified with ofw:GGCCGGCCTCACGG CTTCCCGCCGGCGGTGGCGGCGCAGGATGATGGCACGCTGGTCCCC GTGAAACAGCTGCTCAAC and orev:GCTGAGCAGGTCCAGGA TTTGATTCC. |
| **p*upp3*Δ::P$_{otef}$-mKate2-HA-L2-F2A-Gfp-NLS** | 4198 | NatR | Same as pUMa4197. F2A N-terminally fused to L2 was codon optimized ordered from IDT (Integrated DNA Technologies Inc. Coralville). |
| **p*upp3*Δ::P$_{otef}$-mKate2-HA-L2-Po2A-Gfp-NLS** | 4204 | NatR | Same as pUMa4197. Po2A N-terminally fused to L2 was codon optimized ordered from IDT (Integrated DNA Technologies Inc. Coralville). |
| **p*ip*$^{R}$:P$_{crg}$-*rrm4*-Gfp-L2-P2A-*firefly*-HA** | 4565 | CbxR | Vector for the co-expression of Rrm4 C-terminally fused to Gfp and Firefly luciferase fused to HA tag separated during translation with L2-P2A. L2-P2A was amplified with ofw:TGTACAAACACGGCTT CCCGCCGGCGGTG and orev:CAATTGGGGACCGGGGTTCTCCT CGAC. Firefly was amplified with ofw:CGGGATCCCCCGGGCTG CAGGAATTCGATCCCCAATTGATGGAGGACGCCAAGAA and orev:GCCGGGCGGCCGGCGCGCCGGCCGCTAGATCTTTAGGCGTAGT CGGGCACGTCGTAAGGGTAGAGCGGACCCTGGACGGCGATCTTGCC. |
